# Describing the current status of *Plasmodium falciparum* population structure and drug resistance within mainland Tanzania using molecular inversion probes

**DOI:** 10.1101/2020.05.09.085225

**Authors:** Kara A. Moser, Rashid A. Madebe, Ozkan Aydemir, Mercy G. Chiduo, Celine I. Mandara, Susan F. Rumisha, Frank Chaky, Madeline Denton, Patrick W. Marsh, Robert Verity, Oliver J. Watson, Billy Ngasala, Sigsbert Mkude, Fabrizio Molteni, Ritha Njau, Marian Warsame, Renata Mandike, Abdunoor M. Kabanywanyi, Muhidin K. Mahende, Erasmus Kamugisha, Maimuna Ahmed, Reginald A. Kavishe, George Greer, Chonge A. Kitojo, Erik J. Reaves, Linda Mlunde, Dunstan Bishanga, Ally Mohamed, Jonathan J. Juliano, Deus S. Ishengoma, Jeffrey A. Bailey

**Author notes:** Shared Last Author (Deus S. Ishengoma, Jeffrey A. Bailey). **Corresponding Author Information** Kara A. Moser, PhD, MPH, Institute for Global Health and Infectious Diseases, University of North Carolina Chapel Hill, Bioinformatics, 130 Mason Farm Rd, 2nd Floor, Chapel Hill, NC, USA, 27514, 1-919-966-2537. **Competing interests:** The authors declare that they do not have any competing interests (commercial, or otherwise).

## Abstract

High-throughput *Plasmodium* genomic data is increasingly useful in assessing prevalence of clinically important mutations and malaria transmission patterns. Understanding parasite diversity is important for identification of specific human or parasite populations that can be targeted by control programs, and to monitor the spread of mutations associated with drug resistance. An up-to-date understanding of regional parasite population dynamics is also critical to monitor the impact of control efforts. However, this data is largely absent from high-burden nations in Africa, and to date, no such analysis has been conducted for malaria parasites in Tanzania country-wide. To this end, over 1,000 *P. falciparum* clinical isolates were collected in 2017 from 13 sites in seven administrative regions across Tanzania, and parasites were genotyped at 1,800 variable positions genome-wide using molecular inversion probes. Population structure was detectable among Tanzanian *P. falciparum* parasites, roughly separating parasites from the northern and southern districts and identifying genetically admixed populations in the north. Isolates from geographically close districts were more likely to be genetically related compared to parasites sampled from more distant districts. Known drug resistance mutations were seen at increased frequency in northern districts, and additional variants with undetermined significance for antimalarial resistance also varied by geography. Malaria Indicator Survey (2017) data corresponded with genetic findings, including average region-level complexity-of-infection and malaria prevalence estimates. The parasite populations identified here provide important information on extant spatial patterns of genetic diversity of Tanzanian parasites, to which future surveys of genetic relatedness can be compared.

**SIGNIFICANCE:** Documenting dynamics of malaria parasite genomics in high-transmission settings at scale in sub-Saharan Africa is critical for policy and decision making to support ongoing malaria elimination initiatives. Using molecular inversion probes, we genotyped over 1,000 Tanzanian *Plasmodium falciparum* samples collected country-wide in 2017 at hundreds of variable polymorphic positions across the genome. Frequencies of known drug resistance mutations were higher in northern districts of the country compared to the south. Results also showed a distinct isolation-by-distance pattern (whereby increasing geographic distance was correlated with decreasing genetic relatedness), as well as signals of higher genetic sharing between several southern districts. These results provide, for the first time, a picture of current within-country diversity of Tanzanian *P. falciparum* populations.

## INTRODUCTION

As global malaria morbidity and mortality has decreased over the past two decades, malaria remains entrenched in certain areas. Currently, 70% of the world’s malaria cases occur in 11 countries (1). These “high burden, high impact” nations have recently become the subject of more focused control efforts, shifting how malaria control efforts are implemented and potentially requiring new tools to help monitor and inform public health efforts.

Understanding the parasite dynamics underlying malaria in high transmission regions is critical for ongoing malaria elimination initiatives in sub-Saharan Africa. Monitoring parasite population structure and genomic signatures of selection can offer insights on the impact that interventions have on parasite populations (2), the spread of mutations associated with drug resistance (3), and identify transmission patterns correlating with human movement (4). Recently, high throughput genomic methods have shed light on important geographic (5–7) and temporal (8, 9) patterns of parasite populations, as well as fine-scale genetic relatedness across short distances (10), in Southeast Asia. While these analyses have elucidated within-country dynamics in a region of relatively low transmission, it is unclear how these insights translate to high transmission settings of Africa, home to 10 of the 11 “high burden” countries. Also lacking are thorough baseline genetic characterizations of circulating parasites in these regions, to which subsequent analyses can be compared to. Several studies have managed to identify between-country genetic signatures utilizing whole genome sequence data across sub Saharan Africa (11–14), but a lack of thorough within-country sampling and reliance on whole genome sequencing (which can be cost-prohibitive for large genetic surveys) has limited further exploration of population genetics in Africa.

Recently, molecular inversion probes (MIPs) have been adapted as assays for high-throughput characterization of *Plasmodium falciparum* clinical infections. These assays allow for the rapid characterization of tens to thousands of positions across the genome, and have successfully been used to track the spread of drug resistance mutations (15–17) in African countries and elucidate parasite population structure within the Democratic Republic of Congo (a high burden nation) (18). Expanding MIPs to explore these trends in other high transmission settings in Africa is an efficient way to establish contemporary genetic trends, and can provide ongoing surveillance of circulating parasite populations. Such characterization would provide control programs with baseline and ongoing parasite surveillance data to evaluate public health interventions in high-transmission areas of sub-Saharan Africa that are pivoting to elimination efforts (19, 20).

Tanzania is classified as a high burden, high transmission setting, with over 56 million people (almost the entirety of the population) at risk for malaria (1). However, malaria transmission is heterogeneous across the country (**Figure 1**) (21, 22), with regions of both high and low transmission. Given these patterns, the genetics of underlying parasite populations may be similarly heterogeneous. Additionally, while several studies have investigated patterns of drug resistance markers across the country (23–25), there have been no large-scale analyses of within-Tanzania parasite population structure using genetic markers genome-wide.

**Figure 1:**
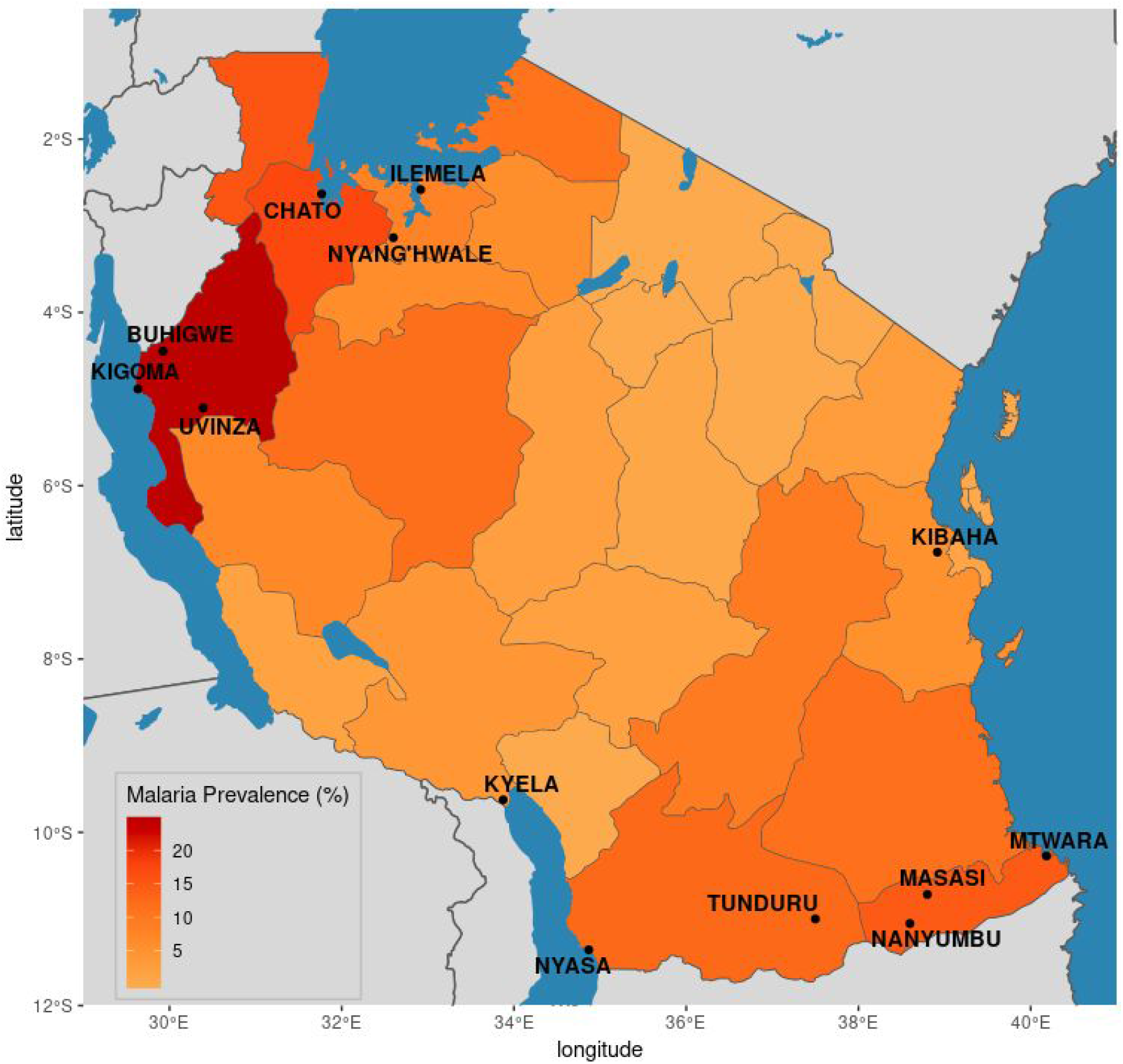
Region-level malaria prevalence in Tanzania. Malaria prevalence in children under five years of age at the regional level, as measured by rapid diagnostic tests (RDTs), was calculated using data from the 2017 Malaria Indicator Survey (MIS). Districts in which sampling occurred are labeled.

In this work, over 1,000 *P. falciparum* infections collected across Tanzania in 2017 were genotyped using MIP panels. Using data generated at ∼1.8k genome-wide positions, as well as known or putative drug resistance loci, a current picture of parasite dynamics within the country is described. Results show that parasite populations differ genetically across the country, and that this differentiation is likely driven by a combination of geographical distance and selection due to past or ongoing antimalarial drug use.

## METHODS

### Sample collection

1,232 samples from 13 districts in 7 regions of Tanzania (**Figure 1**) were available from previously conducted cross-sectional surveys and drug efficacy studies. For the former, 552 samples were collected between July and November 2017 from districts with historically high transmission (22, 26). All individuals in 120 households from two villages selected from each district were tested for malaria, representing both symptomatic and asymptomatic malaria cases in adults and children (27). Parasitemia information from microscopy was available for these subjects. The remaining 680 samples came from two *in vivo* therapeutic efficacy studies (TES) conducted at six national malaria control program (NMCP) sentinel sites in 2017. The first TES assessed the efficacy and safety of artesunate-amodiaquine and dihydroartemisinin-piperaquine in enrolled symptomatic children between 6 months and 10 years of age (28); the second TES evaluated artemether-lumefantrine (Ishengoma *et al.* In preparation). Ethical clearance for the above studies was obtained from the Medical Research Coordinating Committee (MRCC) of the National Institute for Medical Research (NIMR-MRCC) in Tanzania, and analyses utilizing parasite genomes from deidentified samples were deemed nonhuman subjects research by the Institutional Review Boards at Brown University and the University of North Carolina at Chapel Hill.

### Parasite Genotyping and Variant Calling

DNA was extracted from dried blood spots using QIAmp DNA Blood kits per manufacturer’s protocols (Qiagen, Hilden, Germany). Extracted DNA underwent capture and sequencing using two MIP panels that have been previously described (15); the first panel targeted 1,834 single nucleotide polymorphisms (SNPs) across the entire parasite genome, and the second panel targeted SNPs known or suspected to be associated with drug resistance. Libraries were sequenced on an Illumina Nextseq 500 using 150 bp paired end sequencing for the genome-wide panel, and on an Illumina MiSeq using 250 bp paired-end reads for the drug resistance panel at Brown University (MA, USA).

Variant calling was conducted as previously described (15, 18). Briefly, sequences were constructed by stitching together mate-pair reads, and then clustering sequences by 1) their unique molecular identifiers (UMIs) and 2) with the qluster algorithm from SeekDeep (29). SNPs from the resulting sequence were called by pairwise alignment to the reference genome using LastZ (30). Variant calls were subset to the original SNP positions targeted by the MIPs to minimize calls from PCR or sequencing error, and SNPs with poor sequence quality (Illumina phred score < 20) were removed. Non-biallelic SNPs in the genome-wide panel were removed across all samples. After these steps, samples and SNPs were then filtered to remove those that had greater than 90% missing genotypes (across variants or across samples).

### Population Genetic Analyses

To investigate population structure, principal component analysis (PCA) was conducted with the prcomp function in R (v3.6) (31). Discriminatory analysis of principal components (DAPC) was done with the dapc function from the adegenet package (32) (alpha score optimization was used to determine the number of PCs to retain for analysis). Additionally, the program Admixture (33) was used to test *K* values of 1-10 (10 replicates each). The cross-validation (CV) value from the replicate with the highest log-likelihood value for each *K* was plotted, and the *K* with the lowest CV value was chosen as the final *K*. For sample-level analyses using admixture results, samples were assigned to *K* using the within-sample admixture proportion estimates. SNPs associated with geographic regions and *K* subpopulations were explored using the randomForest package in R (34) (ntree=10,000, mtry=200; parameters chosen by observing the out-of-bag (OOB) error over all trees). To investigate the relationship between genetic relatedness and geographic distance (as measured by greater circle distance using district administrative center), the inbreeding_mle function of the mipanalyzer R package was used to calculate the inbreeding coefficient (F) between sample pairs (18). In order to avoid artificially high inbreeding coefficients for samples with high amounts of missing genotypes, this analysis only used samples with less than 50% missing genotypes. For PCA, the within-sample allele frequency (WSAF; allele UMI count divided by total UMI coverage of the site within the sample) was used; for all other analyses, the major allele (represented by > 50% of the UMIs) at heterozygous positions was used.

### Drug resistance analysis

The prevalence of SNPs in known or suspected drug resistance genes was calculated for target SNP positions from the drug-resistance MIP panel at the sample level. If the alternate allele was supported by only a single UMI, or if the WSAF of the alternate allele was less than 1%, the genotype at that position in that sample was considered to only have the reference allele at that position; else the sample was considered positive for the alternate allele. Samples with missing genotypes were excluded from the denominator in prevalence calculations. To explore haplotype information, SNPs that were detected by the same MIP were grouped within each sample, using a unique MIP-microhaplotype ID that is given to each constructed sequence as part of the variant calling pipeline, allowing for the examination of within-sample SNP haplotypes for positions covered by the same probe.

### Epidemiologic Trends

To identify epidemiologic correlations with genetic findings from the above analyses, data from the 2017 Malaria Indicator Survey (MIS) from Tanzania was downloaded from the Demographic Health Survey (DHS) website using the rdhs R package (35). Malaria prevalence and antimalarial use in children under five years of age was calculated using survey weights with the survey package in R (version 3.34). Because the study population presented here included individuals over five years of age, prevalence estimates were also pulled from previously conducted studies within the study districts that included older subjects (36). Complexity of infection (COI) was calculated using the categorical method of THE REAL McCOIL (37) with the genome-wide variants.

## RESULTS

### Population structure

After variant calling and filtering, 742 and 934 samples (from the genome-wide and drug resistance panels, respectively) from 12 districts were retained for analysis (**Table S1**; Methods). High levels of missingness were observed in samples from Nyang’hwale, and were removed (**Figure S1, Figure S2 and Supplemental Text**); after excluding these samples, 737 samples from 12 districts and 1,617 variable biallelic SNPs originally targeted by the MIPs were used to explore population structure in Tanzania. Principal component analysis (PCA) showed separation between the northwest and southern regions of the country, with samples from Kibaha in the east intermingling with samples from both regions (**Figure S3**). Discriminatory PCA (DAPC) analysis using the first 112 components (**Figure S4**) highlighted additional separation of samples by their district-level origin (**Figure 2**), particularly separating samples from the Ilemela district in the north from Kigoma and Buhigwe in the northwest. Admixture analysis identified two main populations (**Figure S5**); one population almost solely represents samples from the northwestern districts, while the other is represented by all districts (**Figure 3A**). Samples from the northwestern districts also had increased admixture, as compared to samples from southern regions.

**Figure 2:**
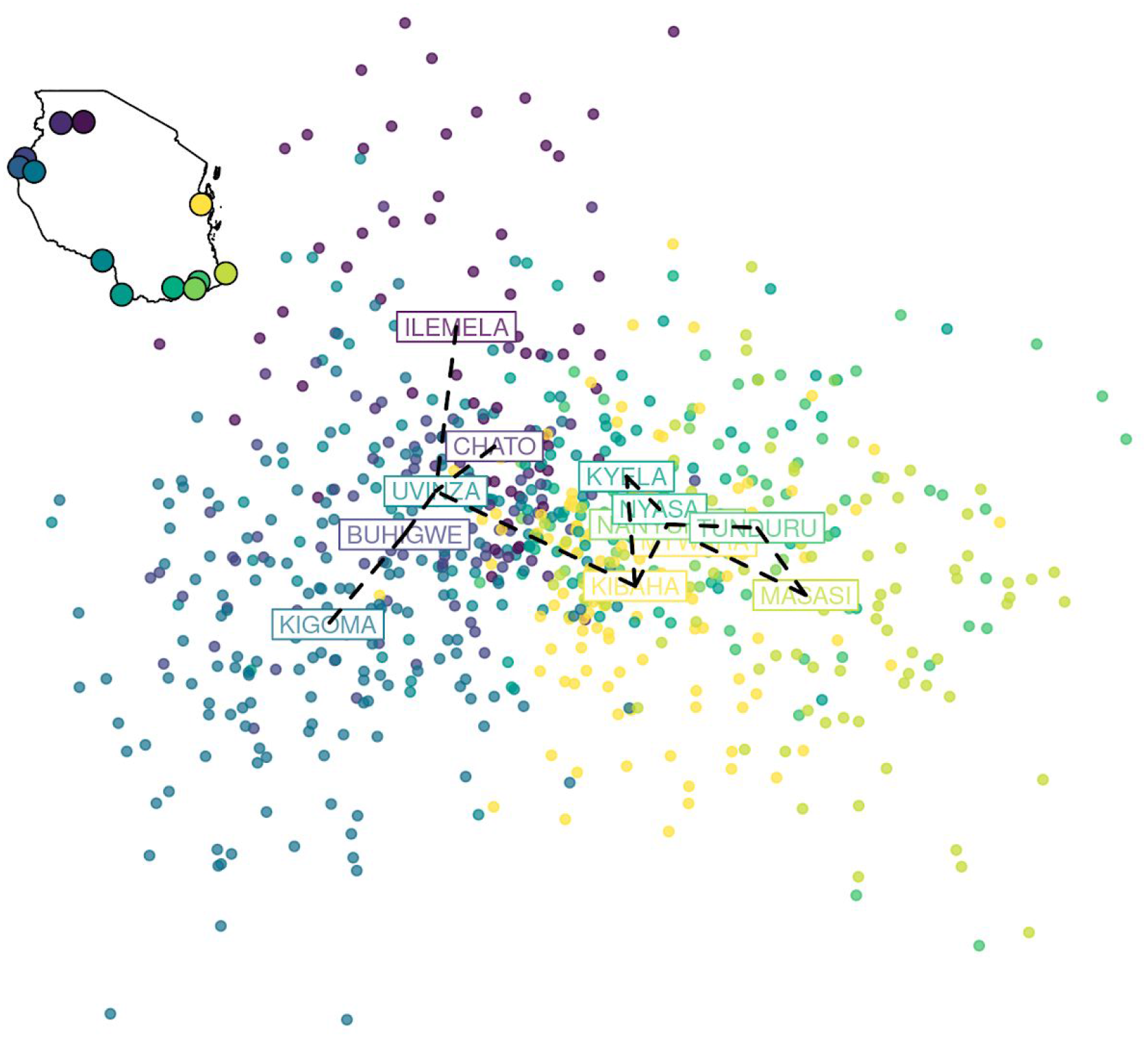
Discriminatory analysis of principal components (DAPC). DAPC analysis using the first 112 components of a principal component analysis (PCA) is shown. Each dot is a sample colored by its geographic origin based on the 12 districts included in the analysis (inset map). A minimum spanning tree, constructed from genetic distances between groups, is overlaid on the plot (black dashed line).

**Figure 3:**
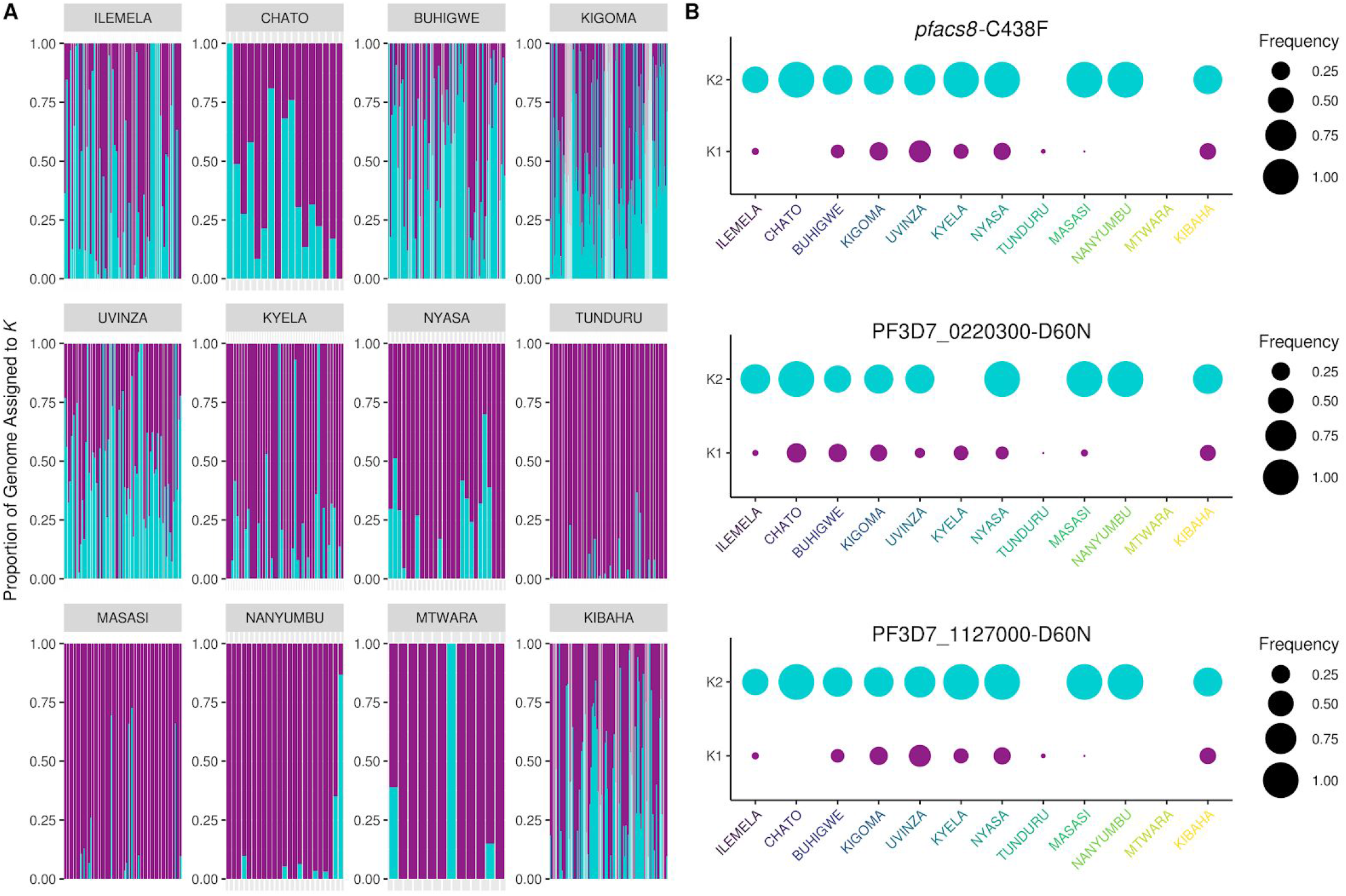
Population structure and associated variants. **A.** Admixture analysis using 737 samples and 1,614 genome-wide SNPs revealed two populations, K1 (purple) and K2 (turquoise) across the 12 districts included in the analysis. **B.** Frequency of key SNPs that were associated with population structure by random forest (Figure S8) or PCA (Figures S9), stratified by subpopulation. Area of the circle represents the frequency of the alternate allele at that position.

As the genetic relatedness of parasites by PCA and DPCA was best represented by a gradient across the sites (versus distinct subpopulations), we estimated genetic relatedness between each sample-pair as the inbreeding coefficient F, which is the probability that any two samples are related by descent, and geographic distance between sample pairs, using only those samples that had at least 50% of targeted SNPs successfully genotyped (n=515). The majority of comparisons showed little to no genetic relatedness (**Figure S6**). However, a small number of samples had inbreeding coefficients exceeding 90%; these highly related samples only occurred between parasites within the same district. After averaging coefficients in bins of increasing geographic distance, an inverse relationship was observed between genetic relatedness and geographic distance (**Figure 4A**). Variation in the amount of genetic sharing also existed by site (**Figure S7**). Sample comparisons from southern district pairs (from within the same district and between different districts) appeared to show slightly more inbreeding than other district-level comparisons (**Figure 4B**, **Figure S7**); additionally, comparisons involving Kibaha did not always track with distance.

**Figure 4:**
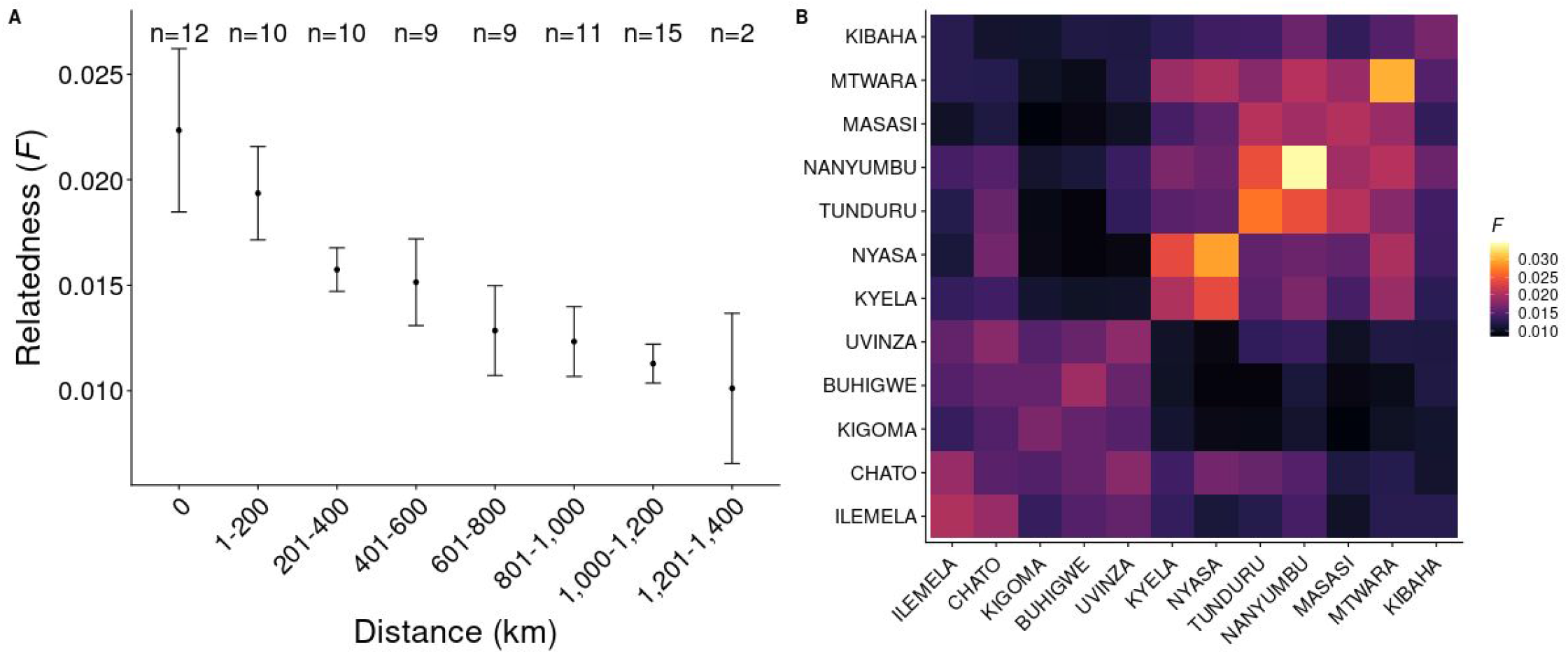
Genetic relatedness by physical/geographic distance. **A.** Relatedness (as measured by the inbreeding coefficient F) binned by geographic distance (km). An average-of-averages approach (average of all district-level average F) was used to avoid district comparisons with more samples contributing more to a bin. Bars represent 95% confidence intervals. The number (n) of district-level pairwise comparisons in each bin is shown at the top of the plot (for example, there are 12 districts included in this analysis, so the number of same-district comparisons in n=12 for bin size 0 km). **B.** Heatmap of averaged F between districts. Districts are arranged by geographic location (north and northwest, to south and southeast, to east).

### Variants associated with population structure

To better understand the genetic drivers of the spatial distribution of parasites, variants associated with the observed population structure patterns were identified using random forest (**Figure S8**). While overall individual variants carried very little predictive power towards classification methods, the top hits on chromosomes 2 and 11 also showed relatively high contributions towards PC1 of the PCA (**Figure S9**). Chromosome 2 hits were in an acyl-CoA synthetase (ACS8; PF3D7_0215300) and a *Plasmodium* exported protein of unknown function (PF3D7_0220300). *pfacs8*-C438F showed particular separation by geography, not only being more prevalent in the northwestern regions of Tanzania, but also occurring more frequently in samples whose genome more likely represented the K2 population identified in the above admixture analysis (**Figure 3B)**(median frequency for K1: 0.04; median frequency for K2: 0.5). Similar results were seen for several SNPs in PF3D7_0220300 (D60N and 146T) and in the chromosome 11 hit PF3D7_1127000 (D60N and K76E). While variants associated with known drug resistance loci were not in among the top hits of the random forest analysis, SNPs in a region upstream of *pfdhps* contributed to PC2 and PC3, while SNPs in a coding region downstream of *pfcrt* (*pfcg2*, a gene often in linkage with *pfcrt* (38)) also contributed to PC3 (**Figure S9**).

### Drug Resistance Patterns in Tanzania

We successfully genotyped and called 934 samples at 32 target SNP positions in 12 genes associated with drug resistance (**Table S2)**. Mutations associated with sulfadoxine-pyrimethamine (SP) resistance in both the *pfdhfr* and *pfdhps* gene were at high frequencies independent of location, such as *pfdhps*-K540**<u>E</u>**. However, other mutations showed differential frequencies across geographic regions, including variants in *pfcrt* and *pfdhps*; the A581**G** mutation in *pfdhps* was observed in the northwest of the country (Kigoma, Buhigwe, Uvinza) but was largely absent from the southern districts (**Figure 5A**); the *pfcrt* mutation K76**T** was also elevated in Chato and Buhigwe (**Figure 5B**). Others, such as A437**G**, had overall high (>50%) frequencies across the country, but still showed fluctuations in frequency, with slightly lower frequencies in the southeastern districts and Kibaha. Haplotypes for these mutations using variants detected on the same MIP probe further highlighted this geographic separation (**Figure S10**). No mutations previously shown to be associated with artemisinin resistance in *pfk13* were observed in any of the districts (**Supplemental Dataset 1**).

**Figure 5:**
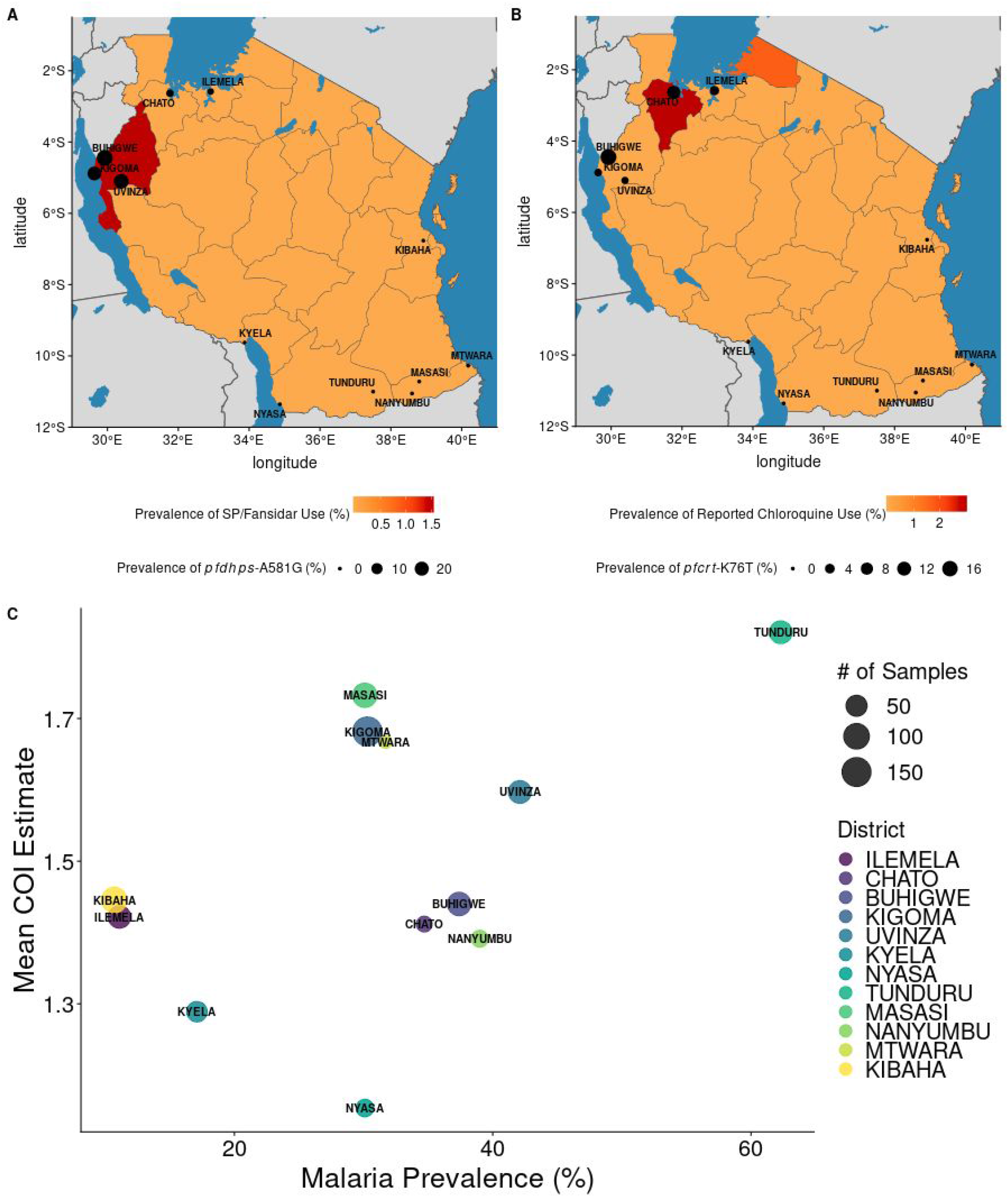
Patterns with antimalarial use and malaria transmission with genetic measures. Data from the 2017 Malaria Indicator Survey (MIS) was assessed for correlation with genetic measures from this study. **A.** District level prevalence of *pfdhps***-** A581**G** mapped on top of region level estimates of SP/Fansidar use among children under five years of age. **B.** District level prevalence of *pfcrt***-** K76**T** relative to region-level estimates of chloroquine use among children under five years of age. **C.** The relationship between *P. falciparum* prevalence, as measured by rapid diagnostic tests (RDTs) in school-age children (36), and average COI estimate, at the district level. Each point is a district, and the size of the district indicates how many samples were in each district. Mutation frequencies for **A.** and **B.** can be found in Supplemental Dataset 1.

### Correlation of Demographic Patterns with Genetic Signals

Higher levels of mutations associated with both SP and chloroquine were observed in northwestern districts (**Supplemental Dataset 1**), despite the replacement of these drugs with artemisinin combination therapies (ACTs) in 2006 (39). To determine if this correlated with ongoing drug use, reported antimalarial use was queried from the 2017 Malaria Indicator Survey data. Reported use of chloroquine, quinine, and SP/Fansidar were all less than 5% (although only ∼20% of participants responded to these queries) (**Figure 5A-B**). Some overlap with regions reporting higher than average use of SP/Fansidar and chloroquine, and districts with >5% prevalence of *pfdhps*-A581G and *pfcrt*-K76T, (**Supplemental Dataset 1**) was observed. Tanzania itself is considered a high-transmission setting; however, malaria prevalence is heterogeneous across the country, with varying levels of prevalence across districts (**Figure 1**). This is also reflected in our genetic data. Complexity of infection (COI) estimates varied by region, with averaged district COI estimates ranging from 1.15 to 1.82. As expected, districts with higher average COI estimates also tended to have higher malaria prevalence (**Figure 5C**), regardless of the source of malaria prevalence estimates (**Figure S11**).

## DISCUSSION

Genomic data of *P. falciparum* infections in malaria endemic regions can provide key information for implementation and control efforts. In addition to monitoring known drug resistance mutations, parasite population genetics are currently being used to characterize important contributions of human mobility to parasite spread (4, 40); such data can allow for the identification of regions of populations that could be targeted by control methods. Genomic data has also been used to assess the impacts of interventions (2), and in the future could be used to monitor for unexpected population shifts after mass drug administration campaigns (the emergence of known and novel drug resistance mutations) and large-scale vaccine feasibility studies (the emergence of vaccine-resistant parasite populations). However, for such studies to be successful, it is important that the background variation in standing parasite populations is well described. The analysis presented here explored patterns of parasite population structure and diversity in combination with drug resistance among parasites in Tanzania. Parasites from these regions are composed of two main populations (north and south). Drug resistance markers are also correlated with geography; several key *pfdhps* and *pfcrt* mutations were observed at higher frequencies in districts from the north and northwest. In addition, patterns of decreasing average genetic relatedness with increasing geographic distance were identified, consistent with predominantly local transmission.

The lack of identification of any sample pairs from different districts with high genetic relatedness indicates that, in this dataset, there is little detectable contribution by very recent human movement to parasite population genetics in this region. However, several signals may still reflect broad patterns in human migration, such as the greater genetic relatedness among samples from Tunduru and surrounding districts. A major highway (A19) directly connects Tunduru to Nanyumbu, Masasi and, eventually, Mtwara. Kyela and Nyasa comparisons also showed greater amounts of inbreeding; both of these districts border Lake Nyasa, and may reflect increased human movement upon waterways. While some southern districts had small sample sizes for this analysis (Nanyumbu, Mtwara Nyasa), high inbreeding was still within and between other southern districts with sample sizes > 40 (Tunduru, Masasi). The increased sharing relative to that seen in regions of similar distance in the north could be due to the north having a more admixed population driving greater genetic distance. However, stratification of the analysis by K1 and K2 subpopulations did not show an expected increase in relatedness (data not shown).

By leveraging previously conducted MIS data, which were collected at a temporally equivalent timeframe as the samples included in the present study, we were able to assess how transmission intensity correlated with COI. Previous studies, including our recent work in the Democratic Republic of the Congo, support that there is a direct relationship between these values, with higher transmission resulting in higher COI (18, 37, 41). In addition, an overall decrease in average COI was observed between the samples from this study (2017) and from Tanzanian samples collected in 2015 that were genotyped in the same manner (18), corresponding to the overall decrease in malaria transmission in Tanzania during that period (42).

The results presented here also confirm earlier work exploring drug resistance mutation patterns in Tanzania, and provide context for how these mutations are occurring. High prevalence of certain mutations across Tanzania, such as *dhps*-K540E mutations, and the higher prevalence of *dhps*-A581G mutations in regions of northern Tanzania compared to southern regions, have been previously described (24, 25). Contextually, high frequencies of both *dhps*-A581G and *pfcrt-K76T* have also been observed in nations bordering the northwestern regions of Tanzania (18), and may be a reflection of historical expansion of drug-resistance parasites in this region of Africa. However, while reported drug use in 2017 was low for SP and chloroquine, it was nonetheless reported in the Tanzanian districts where relatively higher mutation frequencies were observed for SNPs associated with resistance to these antimalarials. Further studies investigating drug use in these regions would be important to tease out any contributions that inappropriate antimalarial use may have on maintaining or further spreading drug resistance mutations among these parasite populations.

Several variants found to associate with population structure here have not previously been shown to contribute to population structure within a country and may be useful for future molecular epidemiology studies tracking parasites within Tanzania. While none of these variants have been definitively tied to antimalarial use, several variants identified here have been reported in studies of resistance. Allele frequencies of *pfacs8* have been shown to differ in Malawi, where SP was used for longer periods of time compared to other African nations (43). In addition, a similar region on chromosome 2 contributed to signals seen in population structure analyses from other African nations, including the Democratic Republic of Congo (18). Finally, PF3D7_0220300 has been reported to be upregulated (along with *pfacs8*) in studies investigating the response of *Plasmodium* parasites to dihydroartemisinin exposure (44). The regions in which these genes fall have also been identified as being under selection across space and time in a study in the Gambia (45). Given drug resistance loci represent some of the strongest signals (14, 18), these variants require future study, particularly in regards to their current distribution across sub-Saharan Africa.

The work presented here highlights the ability of highly-multiplexed targeted sequencing with MIPs to elucidate population structure of *P. falciparum* isolates in areas of high malaria endemicity. Within-country population structure has previously been difficult to observe in sub-Saharan African nations, even with whole genome sequencing. Efficient genotyping with MIPs, coupled with country-wide sampling of large numbers of infections, not only allowed for the identification of population structure across broad geographic regions, but also identified genetic sharing across shorter geographic regions using identity-by-descent based methods. A limitation of this work was the level of missingness of data in many samples with lower parasite densities. This is in part due to MIPS being dependent on a capature step, which compared to standard PCR amplicon sequencing is impacted more by low DNA concentrations or degraded DNA (15). However, this capture step allows for the minimization of errors due to the incorporation of UMIs, and MIPs are more sensitive and less expensive than other methods such as whole genome sequencing. In addition, the lower per-sample cost compared to whole genome sequencing makes these panels amenable to larger population surveys with denser sampling, potentially improving our understanding of parasite populations as sampling strategies can become more rigorous and complex. MIP panels are also adaptable; additional probes can be easily designed and added to existing panels to capture new targets of interest. Despite high sample loss in some districts, patterns of population structure were still identified country-wide, using fewer markers compared to whole genome sequencing, and therefore could be useful tools for monitoring parasite populations over time in this region. As Tanzania continues to make strides in malaria control, including advancing the pre-elimination area on the Zanzibar Archipelago, surveillance of genetic changes of Tanzanian parasite population will be critical for monitoring and optimizing interventions, ensuring elimination success.

## ACKNOWLEDGEMENTS

Authors greatly appreciate the contribution of the research scientists and assistants who took part in the sample collection surveys and TES: William Makunde, Method Segeja, Seth Misago, Daniel Challe, Gasper Lugela, Ezekiel Malecela, Juma Tupa, August Nyaki, Zakayo Nzella, John Masimba, Benson Swai, Fides Mumburi, Francis Chambo, Tilaus Gustav, Masunga Malimi, Juma Akida, Paul Martine, Mwanaidi Mtui, Caroline Minja, Richard Makono, Filbert Francis, Gineson Nkya, Richard Malisa, Raphael Charles, Respigi Kiwango, Lifoba Mosoud, Stumai George, Ali Idris, Ibrahim Materego, Neema Barua, and Michael Makange, Isolide Sylvester, Edwin Liheluka, Cloud Tesha, Emmanuel Chagoha, Happyness Jeremiah, Yahya Derua, Bernard Malongo, Martin Zuakuu, Adam Mgaya, Hashim Ally, Heri Bakari, Alex John, Neema Richard, Rukia Ahmed, Israel Msangi, Shabani Shabani, Joyce Geho, Praygod Swai, Neema Mwangoka, Simon Lugoye, Barnabas Muganda, Hosea Nchana, Claris Ossere, Sofian Mdegela, Moses Sarya, Rosemary Batunika, Jaffari Ramadhani, Charles Chacha, Mary Mwacha, Elinzuu Nicodemu, Juliana Joseph, Stumai George, Mzubwa Paul, Nerbert Mwasote, Caroline Minja, Samwel Bushukatale, Katamapahe Ndimbo, Masudi Ngamaley, Elisha Mg’andile, Hemed Suleiman, Neema Nziku, Peter Kapelanga, Sebastian Kobelo, Eulalia Mwageni, Alex Godwin, Ulrick Mosha, Devotha Mlelwa, Elfrida Kilumbo, and Ruth Mwakyombe. Technical, administrative and logistic support was provided by the finance department at Tanga Centre (Lydia Lugomola, Derick Maira, Joseph Said and Selemani Mandia) and NMCP (Abdallah Kajuna and Judith Kirama); drivers (Seth Nguhu, Francis Mkongo, Saidi Maivaji, Silvester Msamila, Eliwasiri Mmbaga, Thomas Semdoe, Athumani Simba and Ally Mshana) and others (Robert Mhilu, Salome Ngoda, Elfrida Mosha, Haruna Mrisho, Beatrice Semng’indo and Rehema Mtibusa) are highly appreciated. Special appreciation goes to the President’s Office Local Government and Regional Authorities and the village, district and regional authorities, particularly village Leaders, District Executive Directors (DEDs), District Medical Officers (DMOs) and District Malaria Focal Persons (DMFPs) for supporting the research teams during the entire study period. We also thank the Director General of NIMR for granting a permission to publish this paper.

## AUTHOR CONTRIBUTIONS

KAM conducted all analyses, generated figures, and wrote the paper. JAB, DSI, and JJJ conceived the study, provided samples and funding, and wrote the paper. OA, RV and OJW wrote pipelines for variant identification, IBD calculations, and MIS data acquisition, respectively. MD, RAM and PM carried out necessary laboratory analyses for sample preparation and sequencing. AM, MGC, CIM, SFR, FC, SM, FM, RN, MW, RMA, MK, MKM, EK, MA, RAK, GG, CAK, EJR, LM,DB, AL and DSI planned, coordinated and implemented field studies to collect samples and clinical data. All authors reviewed and approved the final version of the manuscript.

## CONFLICTS OF INTEREST & DISCLAIMERS

The authors have no conflict of interest to declare. EJR is affiliated with the CDC; the findings and conclusions in this report are those of the author(s) and do not necessarily represent the official position of the Centers for Disease Control and Prevention/the Agency for Toxic Substances and Disease Registry. RN is a staff member of the World Health Organization and MW is a recently retired staff of the World Health Organization. They alone are responsible for the views expressed in this publication, which do not necessarily represent the decisions, policy or views of the World Health Organization.

## ETHICAL APPROVALS

Ethical clearance for the three studies samples were collected through was obtained from the Medical Research Coordinating Committee (MRCC) of the National Institute for Medical Research (NIMR-MRCC) in Tanzania. This analysis was approved by the IRBs at the University of North Carolina at Chapel Hill. Permission to publish the manuscript was provided by the Director General of NIMR.

## DATA AVAILABILITY

Raw sequencing reads generated through this project have been deposited into the NCBI SRA (Bioprojects PRJNA631258 and PRJNA631263).

