## Supplemental-Text-Tables-Figures for "Describing the current status of *Plasmodium falciparum* population structure and drug resistance within mainland Tanzania using molecular inversion probes"

#### Supplemental Results

<u>*Analysis of missingness in isolates and districts:*</u> Median number of unique molecular identifier (UMI) support was 7 and 2, for the genome-wide and drug resistance panels, respectively. Several locales saw large proportions of samples dropped due to high (>90%) missing genotype calls in the genome-wide panel and drug resistance panel (**Table S1**). While low sample parasitemia was a major contributor to this loss, examining data by experimentally processed 96-well plate showed signs of inefficient capture on several plates; this failure appeared to be largely responsible for increased missingness in certain districts and not necessarily something inherent of parasites from those regions. For example, for the genome-side MIP panel, four out of fourteen plates lost greater than 50% of samples (**Figure S2**), but samples from the same geographic region split between different plates saw higher proportions of missingness on the failed plate than on plates that did not have high failure rates (**Figure S1**). Due to the above plate failure, a high proportion of samples (> 80%) from Chato and Nyang'hwale (both in the northern district of Geita) were lost (**Table S1**). However, 64 (76.5%) samples from Ilemela (also in the northern part of Tanzania) were successfully genotyped, offering representation of parasites from that region in the below analysis. Similar results were seen for the drug resistance MIP panel. After sample processing, we excluded Nyang'hwale from all downstream analyses due to high missigness for all samples from this district; as a sensitivity analysis. Principal component analysis (**Figure S3**) and discriminatory analysis of principal components (**Figure S4, Figure S2**) were repeated with low-missingness Nyang'hwale samples included, with no change in patterns or interpretation of results shown in the figures presented here (data not shown).

### SUPPLEMENTAL TABLES

**Table S1: Success of Genotyping by Site**

| District | Total Collected | 1800 Panel | Drug Resistance Panel |
|---|---|---|---|
| | | < 90% Missing Genotypes | < 90% Missing Genotypes |
| Chato | 115 | 17 (14.8%%) | 95 (82.6%) |
| Nyang’hwale* | 32 | 5 (15.6%) | 15 (46.9%) |
| Buhigwe | 98 | 75 (76.5%) | 76 (77.6%) |
| Kigoma | 197 | 154 (78.2%) | 170 (86.3%) |
| Uvinza | 113 | 67 (59.3%) | 84 (74.3%) |
| Kyela | 101 | 45 (44.6%) | 26 (25.7%) |
| Nyasa | 43 | 26 (60.5%) | 40 (93.0%) |
| Tunduru | 89 | 67 (75.3%) | 80 (89.9%) |
| Masasi | 117 | 86 (73.5%) | 97 (82.9%) |
| Nanyumbu | 39 | 23 (58.9%) | 29 (74.4%) |
| Mtwara | 23 | 12 (52.2%) | 15 (65.2%) |
| Kibaha | 154 | 101 (66.1%) | 132 (85.7%) |
| **Total** | **1,232** | **742** | **949** |

* Excluded from all subsequent analyses

### SUPPLEMENTAL FIGURES

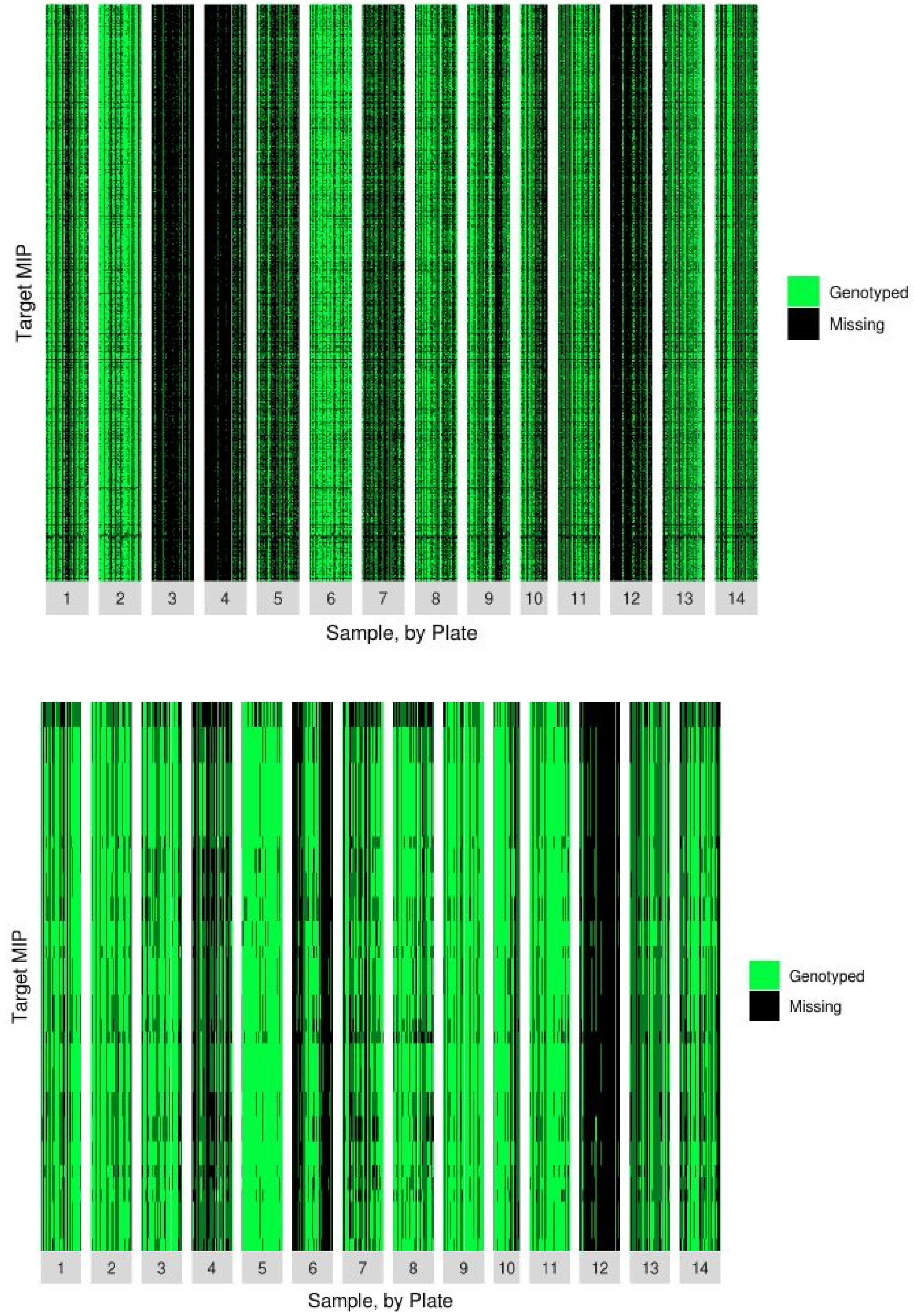


**Figure S1: Relationship between missing genotypes and plate across all samples for the genome-wide MIP panel (top) and drug resistant panel (bottom).** Missing target MIP positions (black) and positions that were successfully genotyped (green) are shown for each sample (column) organized by plate.

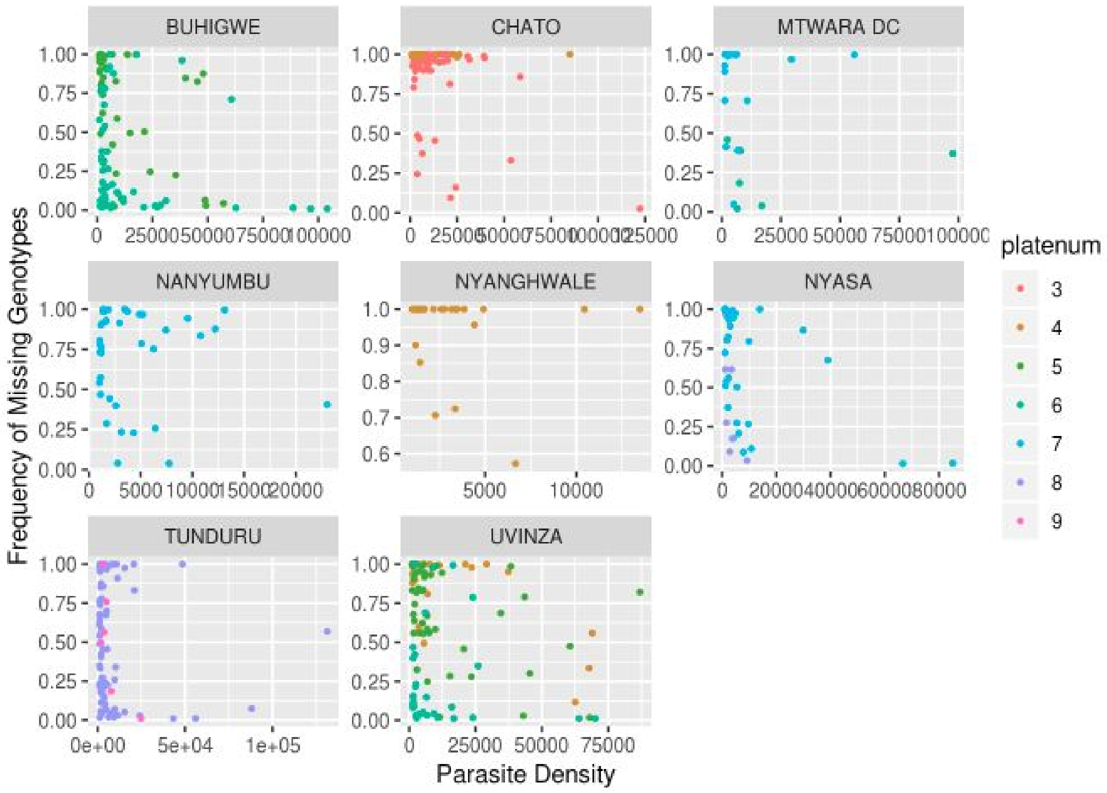


**Figure S2: Relationship between proportion of missing genotypes, parasite density, and plate.** For a subset of samples (n=552), collected as part of the cross sectional study (main manuscript reference 27), parasite density data was available. Each point is a sample, and is plotted by it's parasite density value and the frequency of target MIP sites that failed to be genotyped. Samples are further colored by the plate they were extracted and captured from. (Parasite density data was not available at the time of analysis from the therapeutic efficacy studies.)

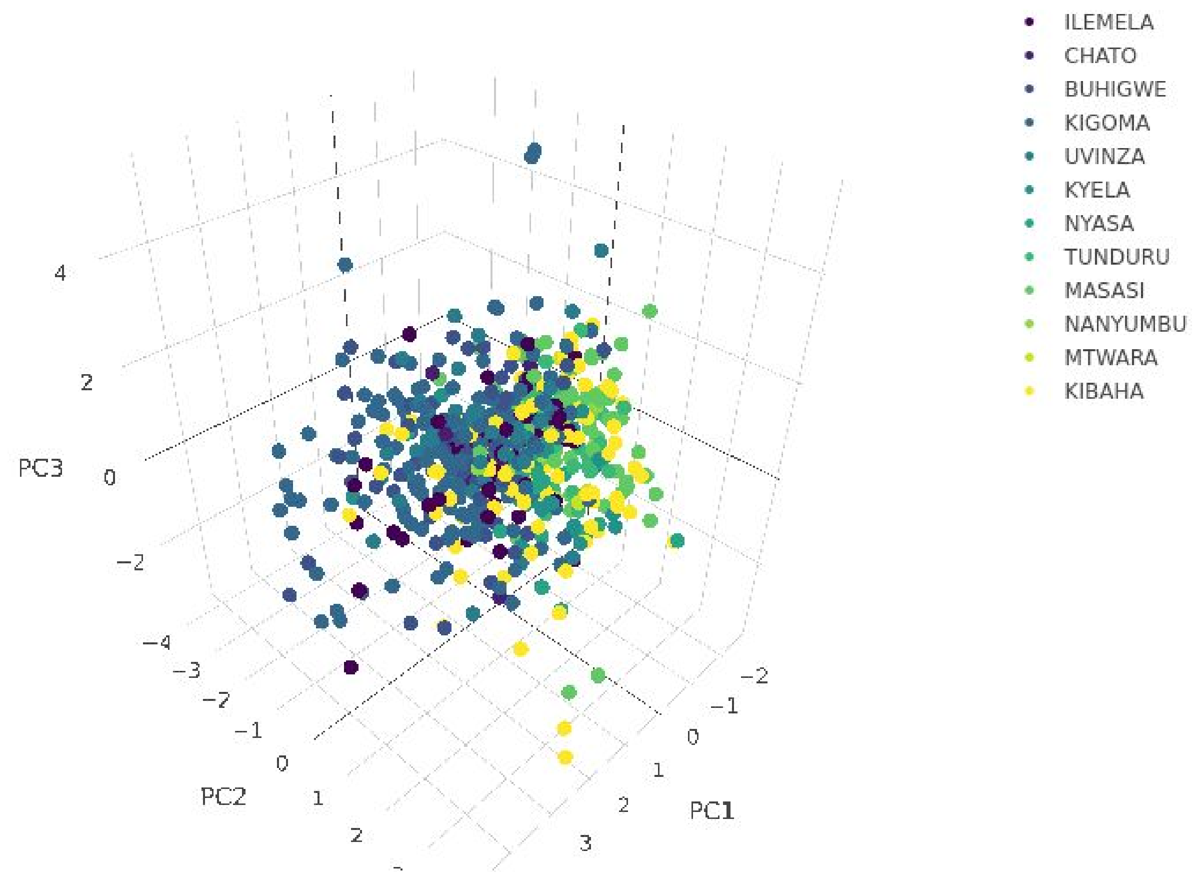


**Figure S3: Principal components analysis (PCA) using genome-wide target SNP positions**. The first three PCA components generated from 737 samples and 1,617 target SNP positions, are shown (explaining 1.37%, 0.92%, and 0.77% of variation in the dataset, respectively). Samples are colored by their district sampling location, and colors are ordered from sites in the north, to the northwest, to the south and southeast, looping back up to Kibaha near Dar es Salaam on the eastern coast.

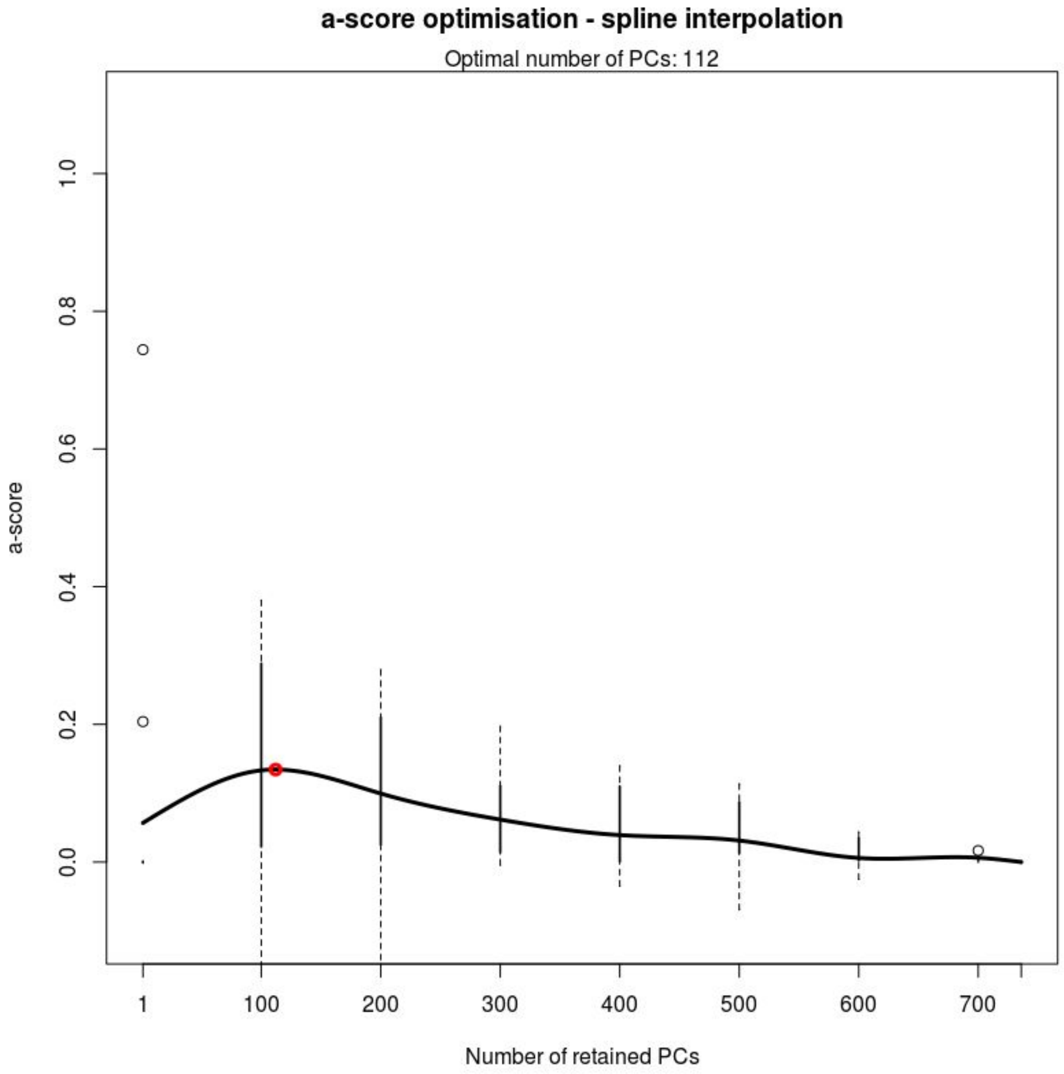


**Figure S4: Ascertaining the number of components to include in the discriminatory analysis of principal components (DAPC).** Alpha-score optimisation shows the optimal number of principal components (112 PCs) to retain for DAPC analysis based on the maximum a-score (red dot).

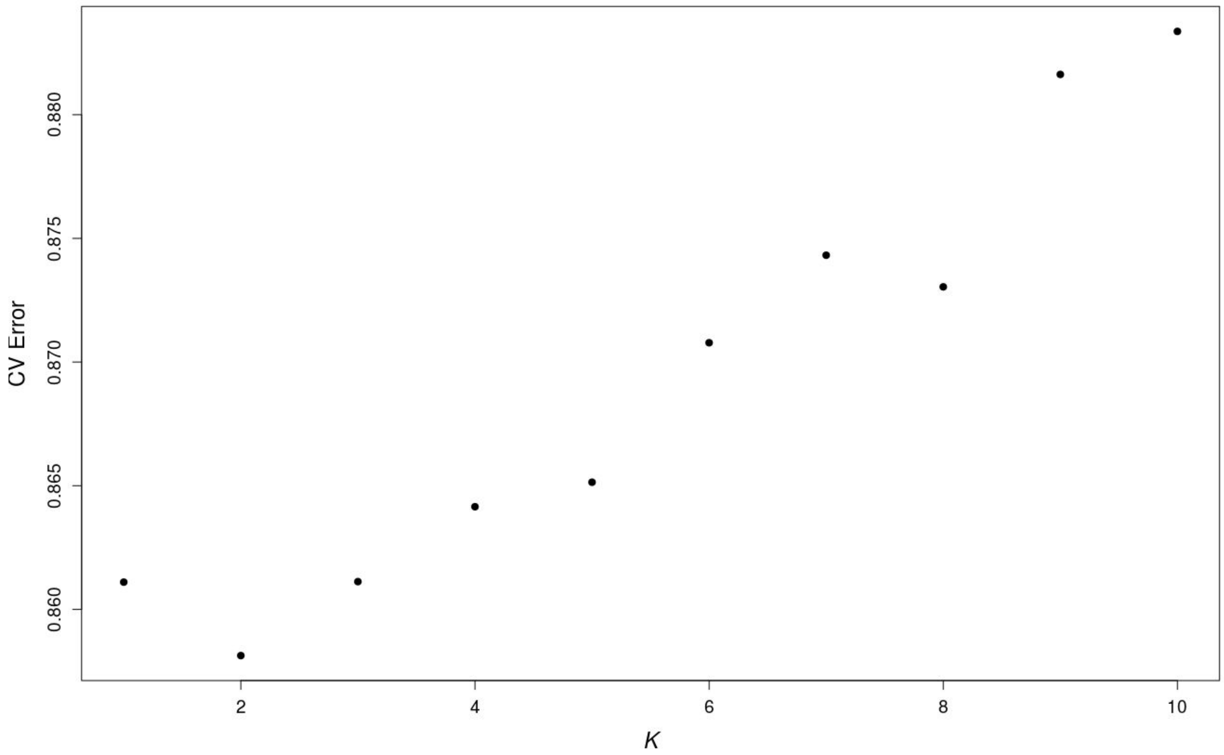


**Figure S5: Identification of best fit population structure.** *K* values of 1 through 10 were tested (x-axis), and the K=2 populations with the lowest cross-validation (CV) error was chosen as the best *K* that supported the given genotype data.

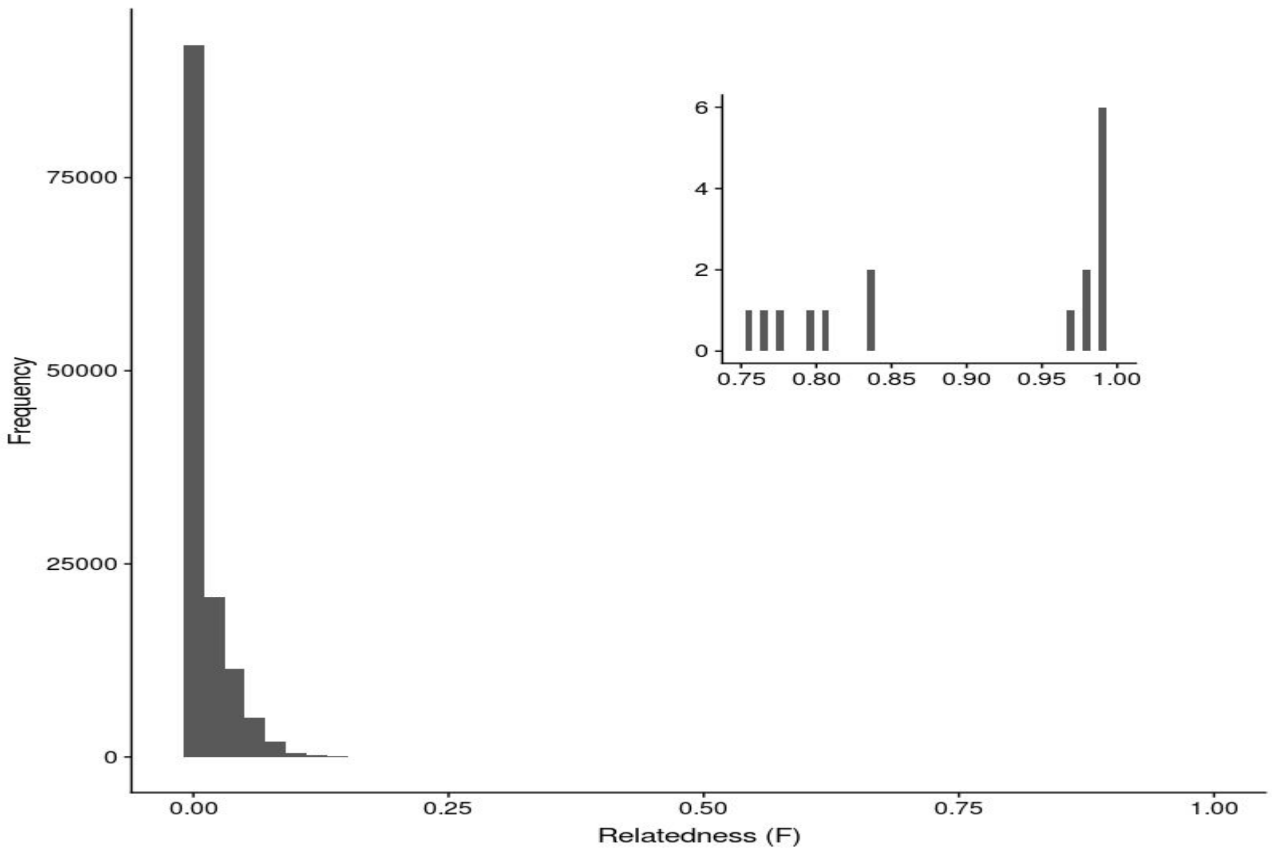


**Figure S6: Inbreeding coefficients (F) between sample pairs.** Each sample was compared to all other samples across the entire sample set (n=515 samples; 132,355 comparisons); the histogram shows the distribution of F coefficients calculated from these sample comparisons. The plot inset shows the smaller number of comparisons that had F coefficient values greater than 0.75, all of which were pairwise comparisons from the same district.

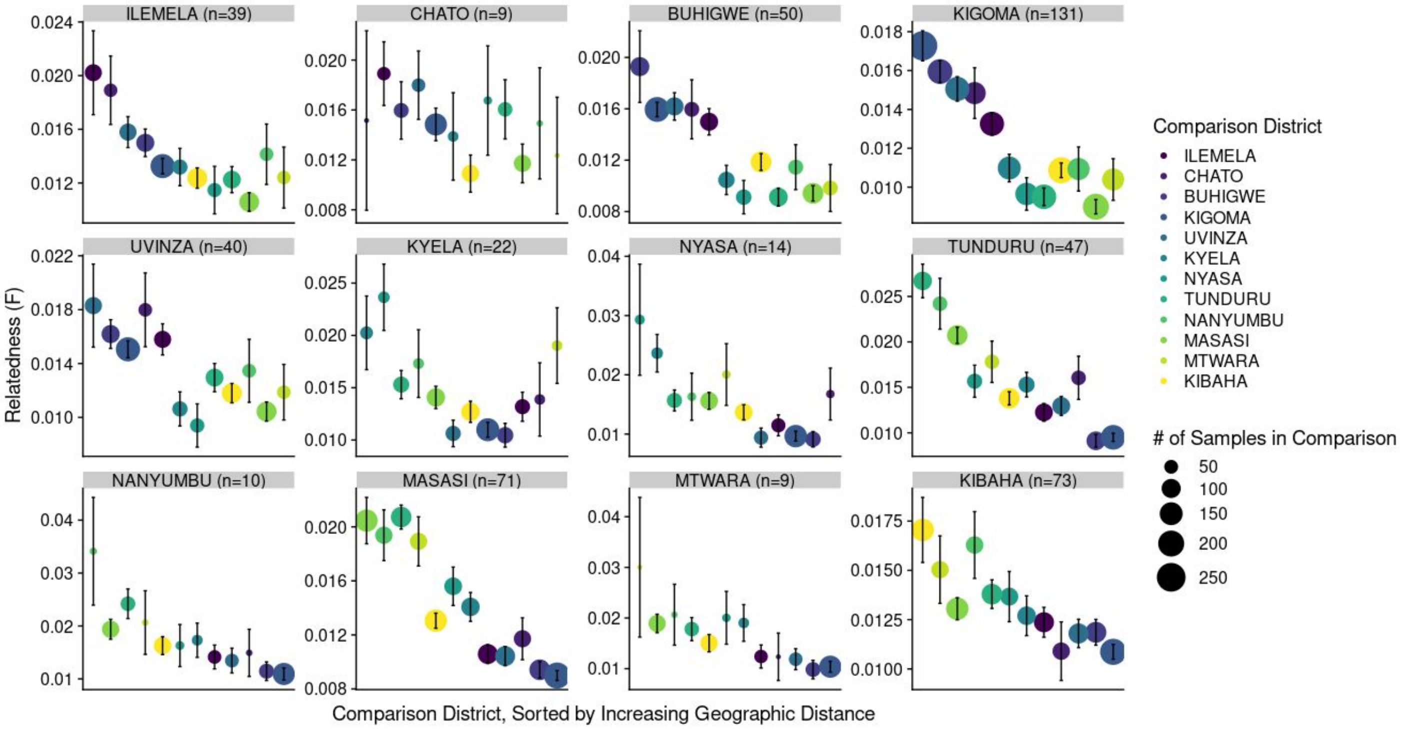


**Figure S7: Geographic distance by genetic relatedness.** Averaged relatedness between any two parasites, as measured by in the inbreeding coefficient (F) (y-axis), decreases as the geographic distance between the districts increases (x-axis). Points are colored by comparison district, and their size is proportional to how many samples went into the pairwise comparisons. Inbreeding coefficient values were averaged across all samples in each comparison group, and 95% confidence intervals are shown.

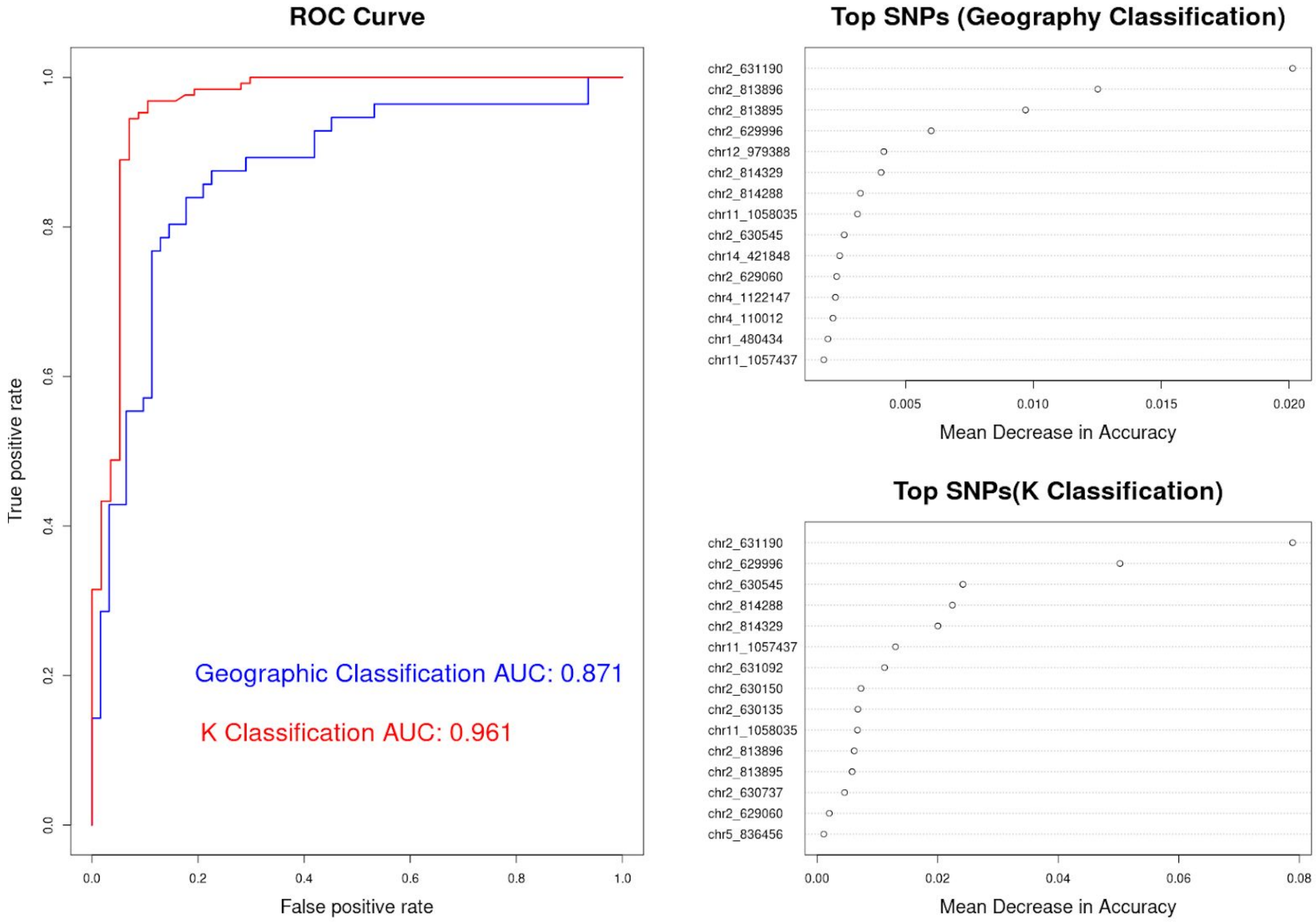


**Figure S8: Results of random forest classification.** Random forests classification models were run on two classifiers: broad geographic region (northern vs southern districts, with Kibaha removed) and sample-level K subpopulation. Balanced training datasets (representing 75% of the data) were used for initial testing, and test data sets (remaining 25% of data) were used to test the predictive accuracy of the classification models. Receiver operating curves (ROC) and area-under-the-curve (AUC) were created by using the model generated from the training dataset to predict classification on the test data set. ROC curves for the models tested on  the test dataset are shown in the left panel. Out-of-bag (OOB) error from the training datasets for classifying by geographic region (21.3%) was much higher than OOB error for classifying by *K* (8.5%), possibly due to the underlying admixture premodinating in samples from the northern districts, and this is also reflected in the AUC values. Top SNPs, sorted by importance, which contribute to the geographic classification model (top right panel) and *K* subpopulation classification model (bottom right).

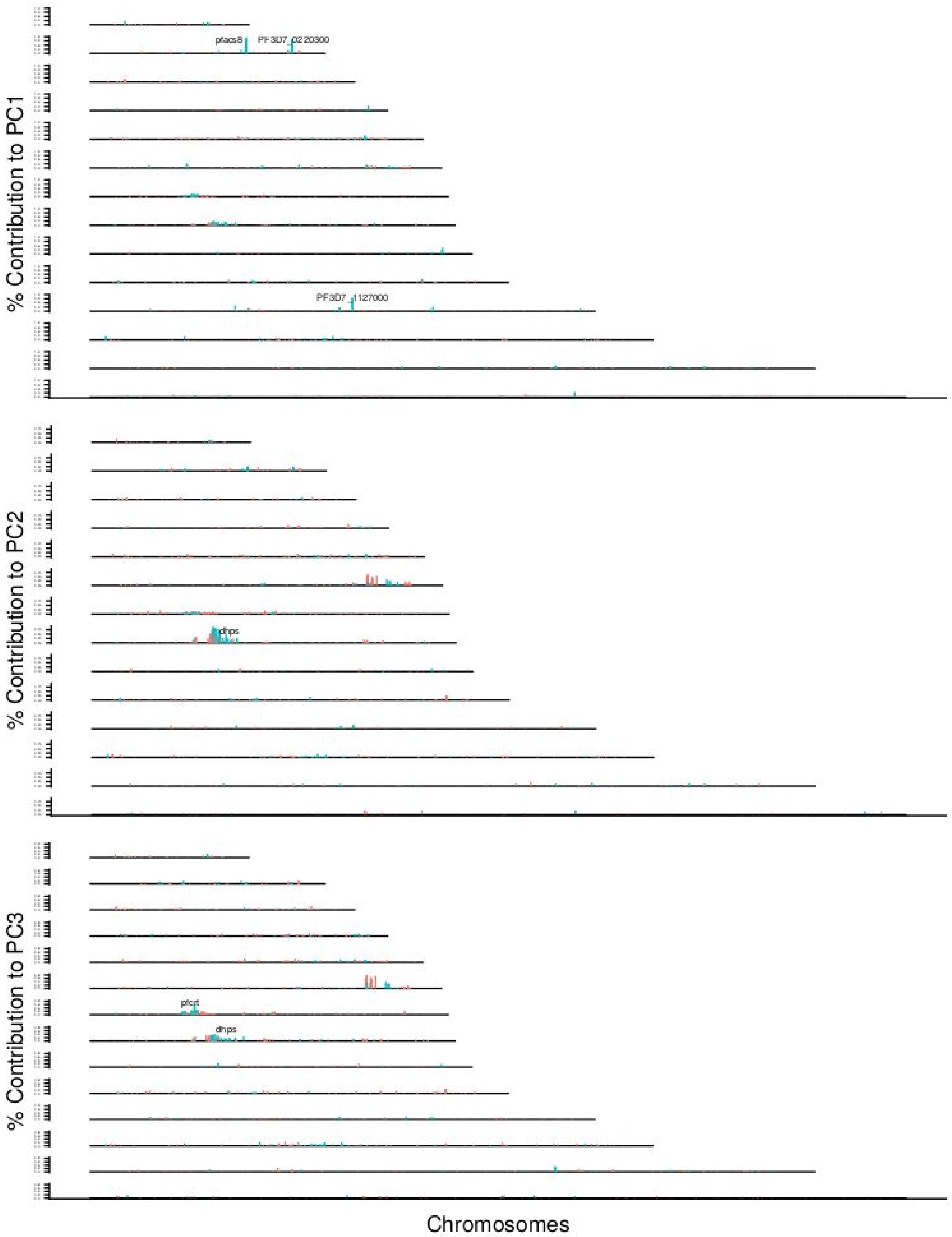

% Contribution to PC1
pfacs8
PF3D7_0220300
PF3D7_1127000
% Contribution to PC2
dhps
% Contribution to PC3
pfcrt
dhps
Chromosomes

**Figure S9: Contribution of individual SNPs to the first three principal components.** Each horizontal line is a nuclear chromosome (ordered from chromosome 1 to chromosome 14); the y-axis for each chromosome is the % contribution to each component. Individual SNPs are colored by coding status (blue: coding; red: non-coding). Select genes have been annotated (labels have been offset in some cases to improve legibility).

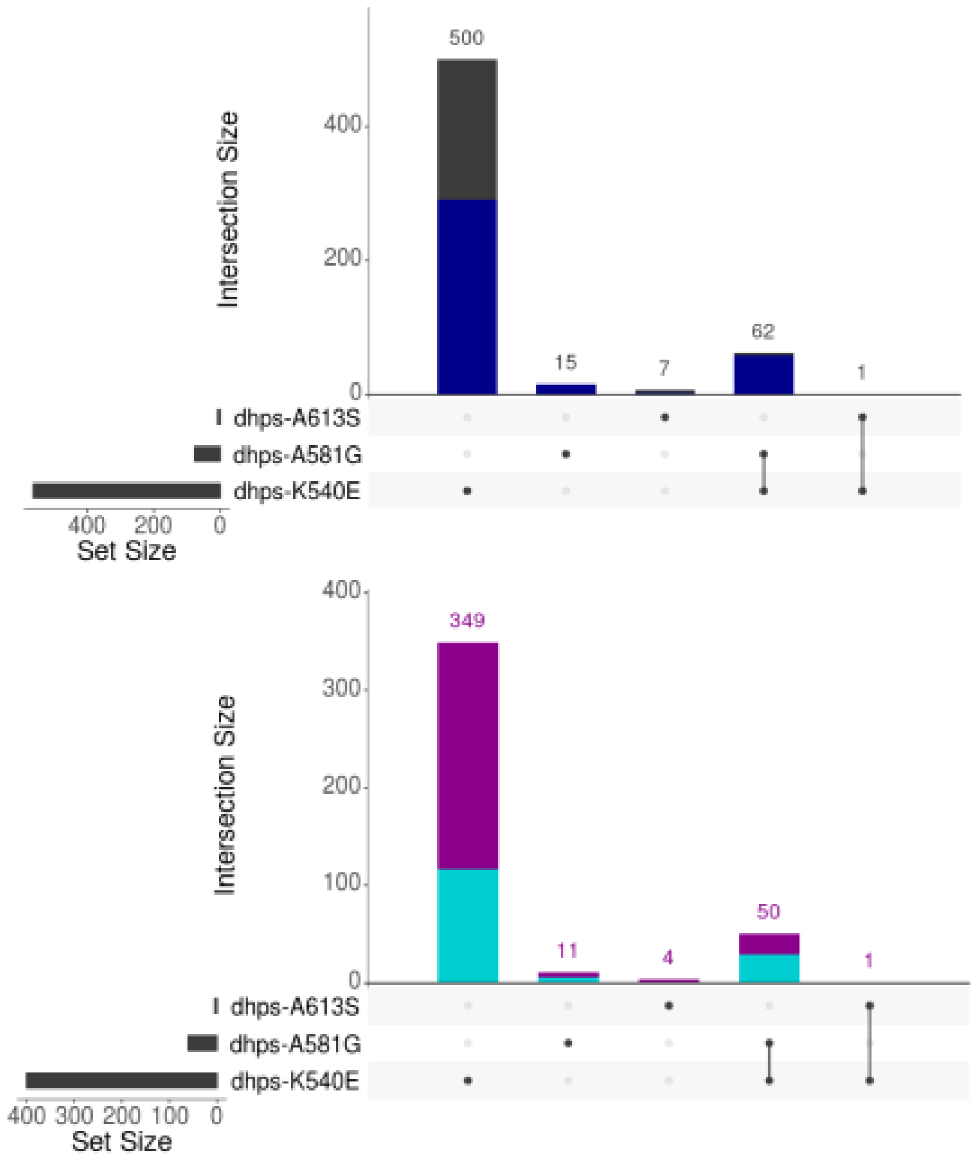


**Figure S10: Haplotypes for *pfdhps* mutations.** Haplotypes for mutations in the *pfdhps* gene within the same MIP probe (K540E, A581G, and A613S) were counted across all samples. **A.** Upset plot showing the frequency of individual and combined mutations. Dark blue represents the proportion of each combination that occured in a sample from the northern geographic region of Tanzania (encompassing Ilemela, Chato, Kigoma, Buhigwe, and Uvinza districts). **B.** The dataset was subset to samples that were also successfully genotyped with the genome-wide MIP panel and assigned into one of the two *K* subpopulations identified in the admixture analysis (Figure 2A). The Upset plot is now colored by a sample's assignment to either K1 (purple) or K2 (turquoise). As MIP information allows the reconstruction of multiple haplotypes within an infection, totals may not always match those in Table S2.

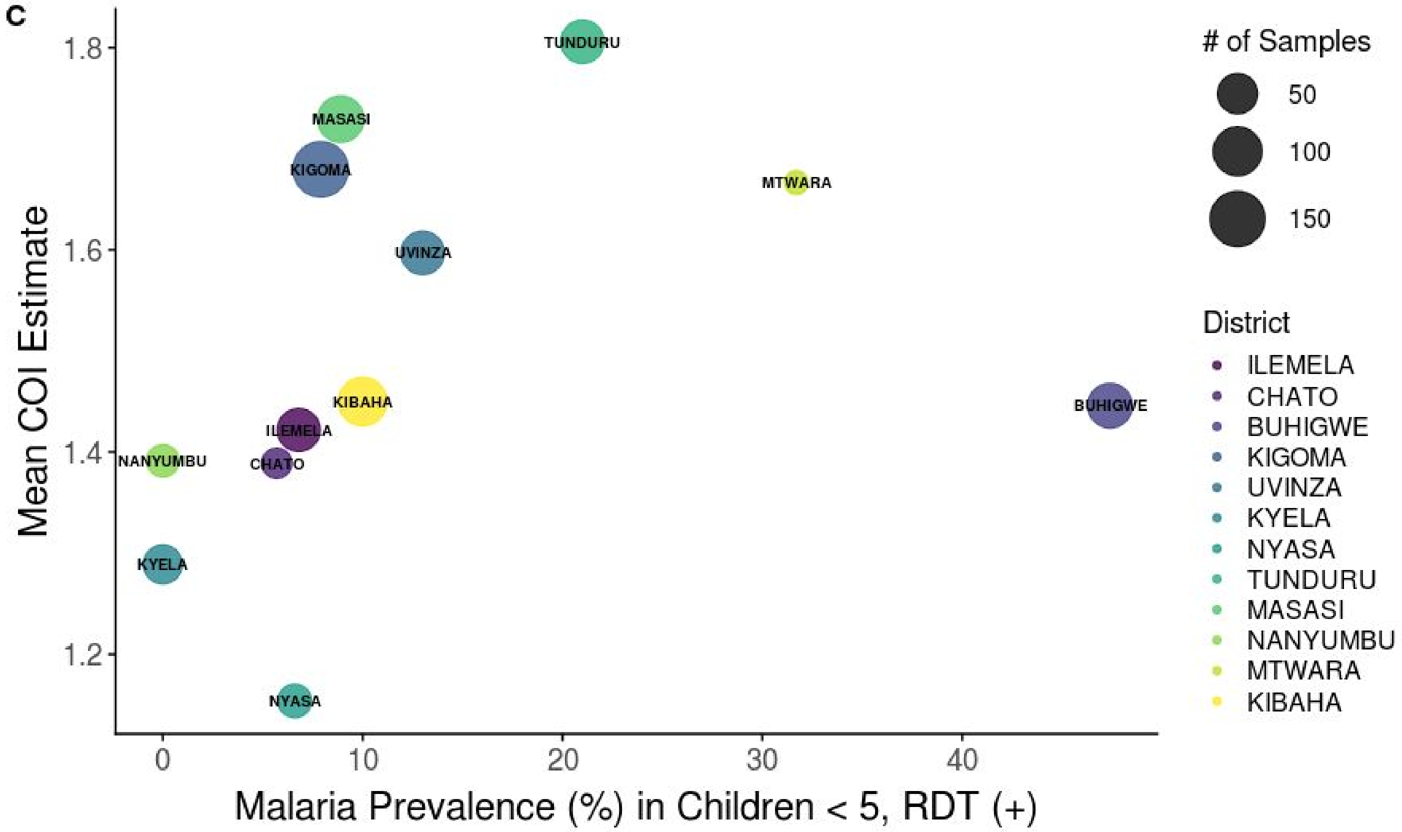


**Figure S11: Correlation of complexity of infection and district level malaria prevalence using MIS 2017 data**. The relationship between *P. falciparum* prevalence from the MIS 2017, as measured by rapid diagnostic tests (RDTs) in children six months to five years of age, and average COI estimate, at the district level. Each point is a district, and the size of the district indicates how many samples were in each district. District level estimates of prevalence were calculated using the MIS 2017 data with the survey package in R.
